# Antiviral Drug Screen of Kinase inhibitors Identifies Cellular Signaling Pathways Critical for SARS-CoV-2 Replication

**DOI:** 10.1101/2020.06.24.150326

**Authors:** Gustavo Garcia, Arun Sharma, Arunachalam Ramaiah, Chandani Sen, Donald Kohn, Brigitte Gomperts, Clive N. Svendsen, Robert D Damoiseaux, Vaithilingaraja Arumugaswami

## Abstract

Emergence of a highly contagious novel coronavirus, SARS-CoV-2 that causes COVID-19, has precipitated the current global health crisis with over 479,000 deaths and more than 9.3 million confirmed cases. Currently, our knowledge of the mechanisms of COVID-19 disease pathogenesis is very limited which has hampered attempts to develop targeted antiviral strategies. Therefore, we urgently need an effective therapy for this unmet medical need. Viruses hijack and dysregulate cellular machineries in order for them to replicate and infect more cells. Thus, identifying and targeting dysregulated signaling pathways that have been taken over by viruses is one strategy for developing an effective antiviral therapy. We have developed a high-throughput drug screening system to identify potential antiviral drugs targeting SARS-CoV-2. We utilized a small molecule library of 430 protein kinase inhibitors, which are in various stages of clinical trials. Most of the tested kinase antagonists are ATP competitive inhibitors, a class of nucleoside analogs, which have been shown to have potent antiviral activity. From the primary screen, we have identified 34 compounds capable of inhibiting viral cytopathic effect in epithelial cells. Network of drug and protein relations showed that these compounds specifically targeted a limited number of cellular kinases. More importantly, we have identified mTOR-PI3K-AKT, ABL-BCR/MAPK, and DNA-Damage Response (DDR) pathways as key cellular signaling pathways critical for SARS-CoV-2 infection. Subsequently, a secondary screen confirmed compounds such as Berzosertib (VE-822), Vistusertib (AZD2014), and Nilotinib with anti SARS-CoV-2 activity. Finally, we found that Berzosertib, an ATR kinase inhibitor in the DDR pathway, demonstrated potent antiviral activity in a human epithelial cell line and human induced pluripotent stem cell (hIPSC)-derived cardiomyocytes. These inhibitors are already in clinical trials of phase 2 or 3 for cancer treatment, and can be repurposed as promising drug candidates for a host-directed therapy of SARS-CoV-2 infection. In conclusion, we have identified small molecule inhibitors exhibiting anti SARS-CoV-2 activity by blocking key cellular kinases, which gives insight on important mechanism of host-pathogen interaction. These compounds can be further evaluated for the treatment of COVID-19 patients following additional in vivo safety and efficacy studies.

**Disclosures:** None declared.

## INTRODUCTION

The current pandemic is caused by a newly discovered coronavirus, severe acute respiratory syndrome-related coronavirus 2 (SARS-CoV-2). As of today, the disease has spread to 213 countries or territories, and the number of COVID-19 cases has surpassed 9.3 million globally, with over 479,000 deaths^1,2^. SARS-CoV-2 is an enveloped single positive-sense RNA virus, which codes a large non-structural polyprotein ORF1ab, four structural proteins and five accessory proteins^3^. The virus enters into host cell by binding its transmembrane spike glycoprotein (S protein) to the cellular membrane angiotensin converting enzyme 2 (ACE2) receptor^4^. ACE2 is expressed in various organs, including lung, heart, kidney, and small intestine^4^. A recent study showed a gradient of ACE2 expression in the respiratory tract with the highest levels in the nose and decreasing expression in the lower respiratory tract^5^. In the proximal airway, SARS-CoV-2 infected ciliated cells, while type 2 alveolar cells (AT2) were found to be infected in the distal airway^5^. The major causes of morbidity and mortality from COVID-19 are acute lung injury with diffuse alveolar damage resulting in acute respiratory distress syndrome (ARDS)^6^. This results from viral replication in lung epithelial cells causing cell injury, a vigorous immune response, respiratory failure, and death. Moreover, there have been additional reports of patients exhibiting acute kidney injury^7-9^, vascular inflammation (endotheliitis), and cardiac complications^10^. Underlying cardiac ailments, diabetes, and obesity are linked to with increased risk of mortality^11,12^. Developing a therapeutic that prevents viral replication is likely to significantly reduce the severity of COVID-19 disease in affected individuals.

Our strategy for developing COVID-19 treatment is based on two main facts about the disease: 1) all patients presenting with symptoms have been infected with SARS-CoV-2 and the virus has gained entry into the airway cells and spread systematically. Given the high affinity and potential avidity in the viral Spike-cellular ACE2 receptor interaction, targeting the viral receptor to prevent cell entry may not a viable strategy to treat patients with this infection, and 2) viruses are dependent on cellular proteins for each step of their life cycle and they hijack many of the host cell factors for their replication. Moreover, these host proteins are not subjected to evolutionary pressure at the short duration of acute infection, therefore there is limited chance of emergence of drug resistant viral mutants. On the other hand, RNA viruses mutate during each round of genome replication due to error prone nature of viral RNA-dependent RNA-polymerase (RdRp) and develop resistance to direct acting antivirals agents (DAA’s). For Influenza A, resistance to adamantanes quickly increased from 1.9% in the 2004 influenza season to 92.3% in 2005-2006 season^13,14^. Viruses take over a large number of host kinases at distinct steps of their life cycle^15- 18^, thus the kinases represent attractive targets for broad-spectrum therapy. These findings, combined with the development and approval of a large number of kinase inhibitors for the treatment of cancer^19^ and inflammatory conditions^20^ have sparked efforts aimed to determine the therapeutic potential of such drugs to combat viral infections.

Currently approved antiviral drugs treat fewer than ten viral infections. The majority of these drugs are direct-acting antivirals that target proteins encoded by individual viruses. As such, this approach provides a narrow spectrum of coverage and therefore cannot address the large clinical need. The high average cost (over two billion dollars) and long timeline (8–12 years) to develop a new drug^21^, further limit the scalability of the DAA approach to drug development, particularly with respect to emerging viruses. This approach is therefore not feasible for the short-term development of a cure that is specific for SARS-CoV-2. The screening of approved drugs to identify therapeutics for drug repurposing is therefore a valid and more universal approach, and several approved drugs have been identified as having activity against many viral diseases^22-26^. Previous drug screening of FDA approved compounds for SARS-CoV, MERS-CoV, and SARS-CoV-2 demonstrated the efficiency of screening approved or clinically developed drugs for identification of potential therapeutic options for emerging viral diseases, which also provided an expedited approach for supporting off-label use of approved therapeutics ^26,27^.

The objectives of this study are: 1) to identify small molecule kinase inhibitors with anti SARS-CoV-2 activity, and 2) to understand the mechanism of host-pathogen interactions by defining the cellular signaling pathways critical for SARS-CoV-2 replication.

## RESULTS

### Infectious SARS-CoV-2 cell culture system

To identify antiviral compounds that could be used for COVID-19 treatment, we established a SARS-CoV-2 infectious cell culture system and virological assays using Vero-E6 cells. SARS-CoV-2, Isolate USA-WA1/2020, was obtained from BEI Resources of National Institute of Allergy and Infectious Diseases (NIAID) and all studies involving live virus was conducted in UCLA BSL3 high-containment facility. SARS-CoV-2 was passaged once in Vero-E6 cells and viral stocks were aliquoted and stored at -80°C. Virus titer was measured in Vero-E6 cells by TCID50 assay. Striking cytopathic effect (CPE) was observed in SARS-CoV-2 infected cells (Figure 1A), indicating viral replication and associated cell injury. At 48 hours post infection (hpi), viral infection was examined by immunofluorescent (IFA) analysis using SARS-CoV Spike (S) antibody. Spike protein was detected in the cytoplasm of the infected cells, revealing presence of viral infection (Figure 1B). We also demonstrated that the drugs, hydroxychloroquine (HQ; 10 µM), a known endosomal acidification inhibitor, as well as interferon-β effectively blocked SARS-CoV-2 infection^28^ (Figure 1C). Therefore, we used this platform has been used for the subsequent drug screening studies.

**Figure 1.**
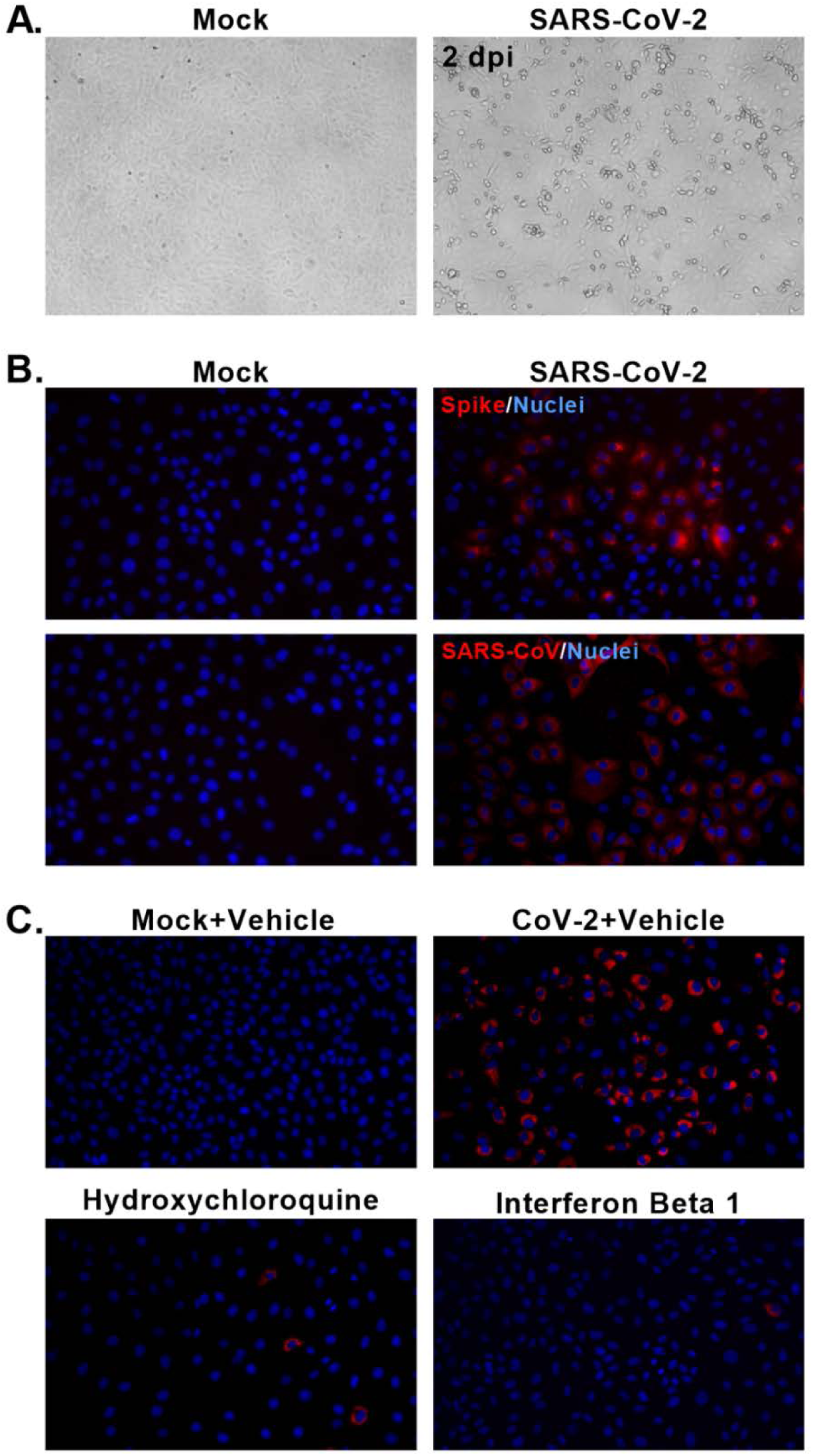
Infectious SARS-CoV-2 cell culture system. (A) Bright field images of Vero-E6 cells infected with SARS-CoV-2 virus. Viral cytopathic effects are noted in the infected culture. 10X magnification. (B) IFA images of infected cells (48 hpi) stained for SARS-CoV-2. Mouse monoclonal antibody (MS Ab) targeting Spike and a guinea pig polyclonal SARS-CoV antibody (GP) were used. Mock infected cells were included as negative control. 20X magnification. (C) IFA images show compounds with antiviral activity. Images depict that hydroxychloroquine (HQ) drug efficiently inhibited SARS-CoV-2 infection. The mock and infected cells were immunostained with dsRNA antibody, which recognizes double stranded genomic RNA generated during viral replication. 20X magnification. Representative data from three or more independent experiments are presented.

### High throughput drug screen of Kinase inhibitors

Next, we set out to evaluate the antiviral property of cellular protein kinase inhibitors. A step by step workflow of our drug screen is provided in Figure 2A. The drug compound library was selected to broadly cover kinase inhibitors and the screening concentration was identified that would have optimal activity and minimal toxicity. We tested a set of 430 kinase inhibitors that have typically been tested for oncologic and immunologic indications and which also have Phase 1/2/3 data available (Supplementary Table 1). However, data for these compounds are not available on SARS-CoV-2. Since this kinase inhibitor library is targeting cancer indications, we decided to avoid using human lung cancer epithelial cell lines. Compounds were formulated into DMSO and pre-plated into media at a 2x concentration (final drug concentration 250 nM). Compounds were added to the Vero-E6 cells in the BSL-3 laboratory followed by the SARS-CoV-2 at a Multiplicity of Infection (MOI) 0.1. After the 48-hour incubation at 37°C, 5% CO2, viral CPE was scored and imaged (Figure 3). The compounds that prevented the viral CPE were identified (Table 1 and Supplementary Figure 1) and subjected to pathway analysis. The drug-cellular protein interaction network was created by mining the collection of 34 hit compounds against the STITCH database in an unbiased fashion, using the standard and unmodified settings^29^. The hit compounds targeted just a few selected kinases, such as mTOR, AKT, PI3K, SRC, ABL, and ATR and a very limited set of pathways, mTOR-PI3K-AKT, ABL-BCR/MAPK, and DNA-Damage Response (DDR) (Figure 2B), suggesting the specific nature of the identified antiviral agents.

**Table 1.**
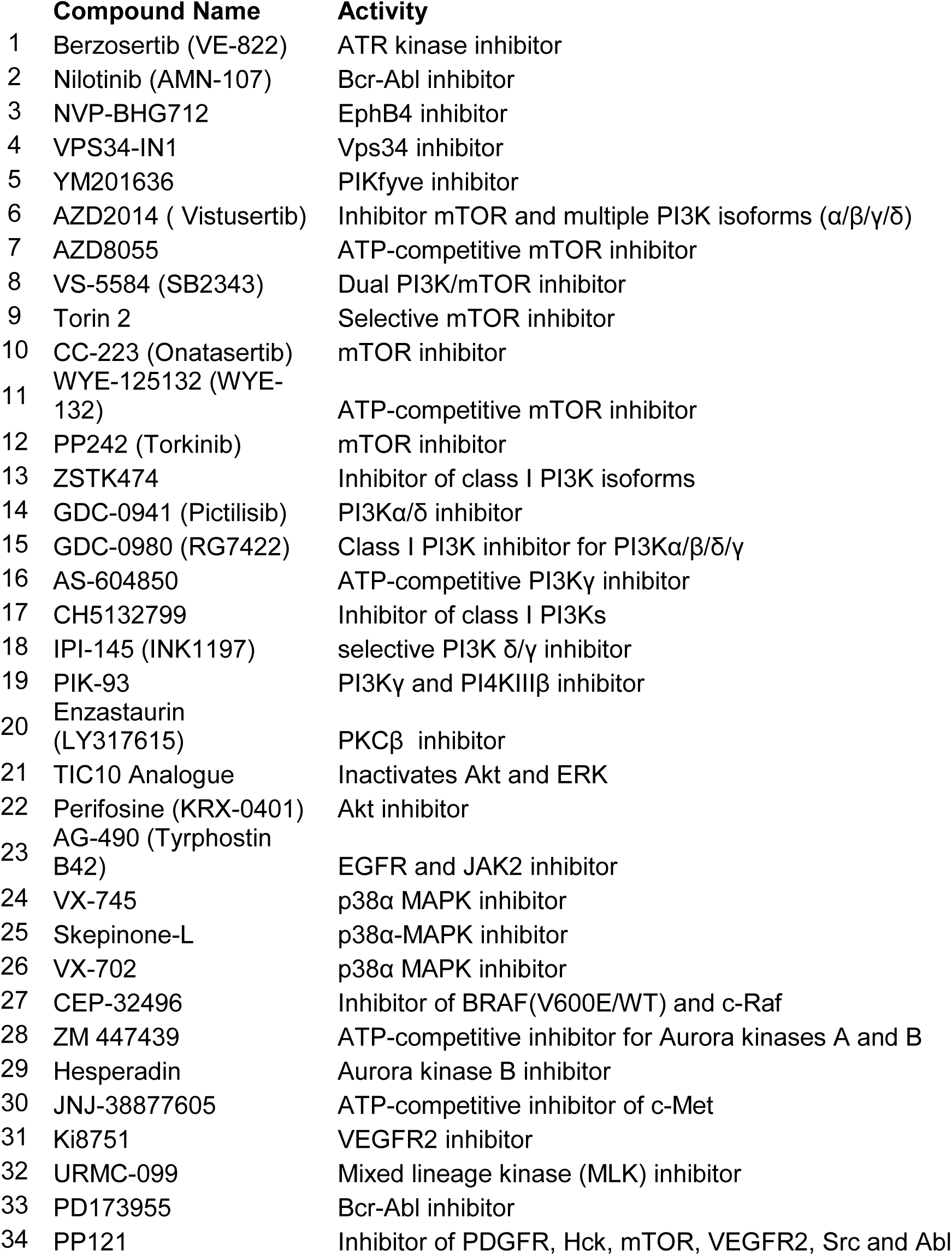
Drugs compounds selected from primary screen having antiviral activity at 250 nM concentration.

**Figure 2.**
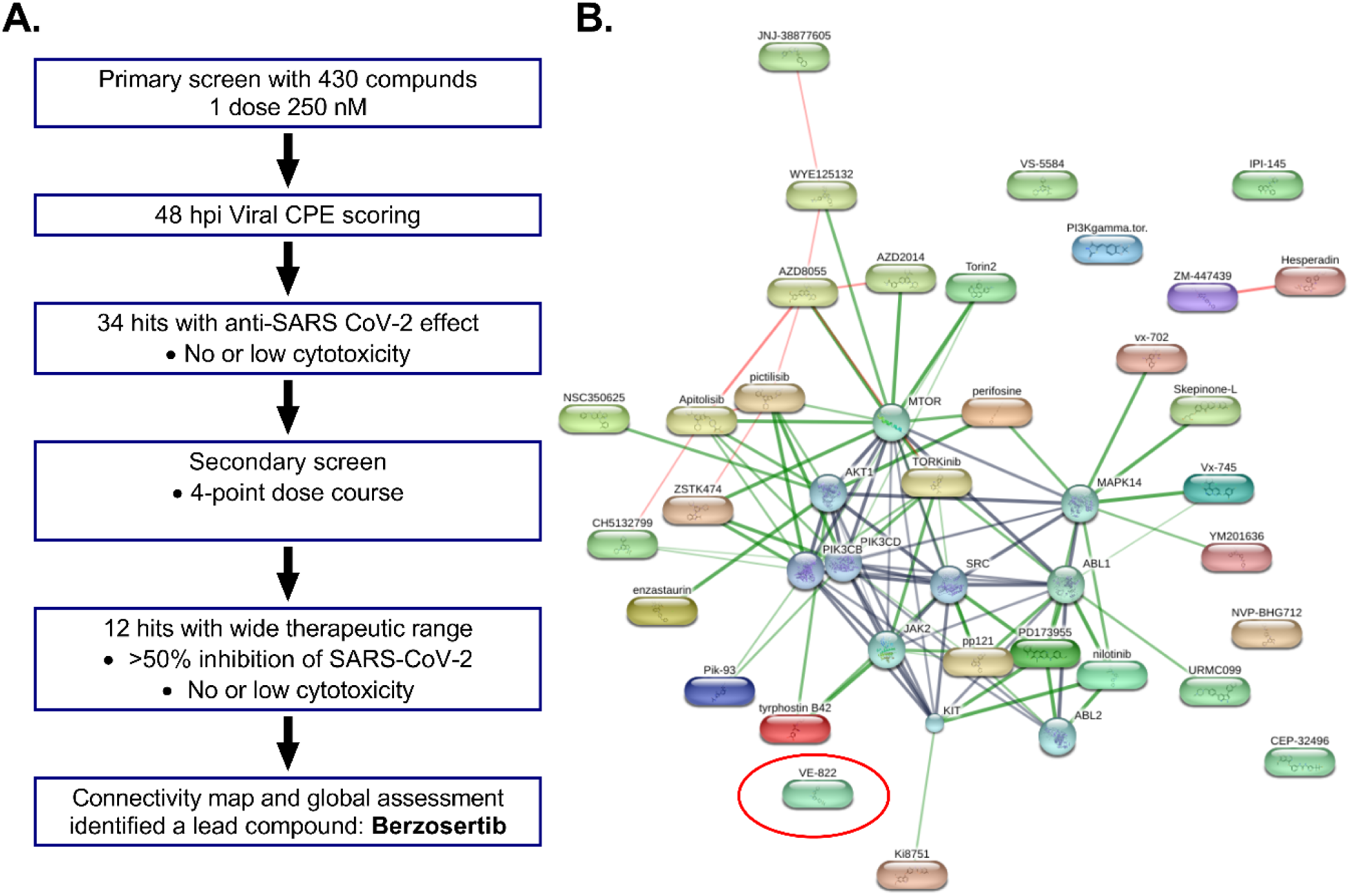
Workflow of high-throughput screening and drug-target kinase connectivity network. (A) Workflow of high-content screen is shown. (B) Connectivity map of 34 drug hits from the primary screen is illustrated. The graphical representation shown is the confidence view in which stronger associations are represented by thicker lines, protein-protein interactions are shown in grey, chemical-protein interactions in green and interactions between chemicals in red. Round shapes represent proteins and oval shapes indicate hit compounds from the primary screen. The analysis indicated a protein-protein interaction enrichment score of 0.0026, which is statistically significant. VE-822 (Berzosertib) is circled in red.

**Figure 3.**
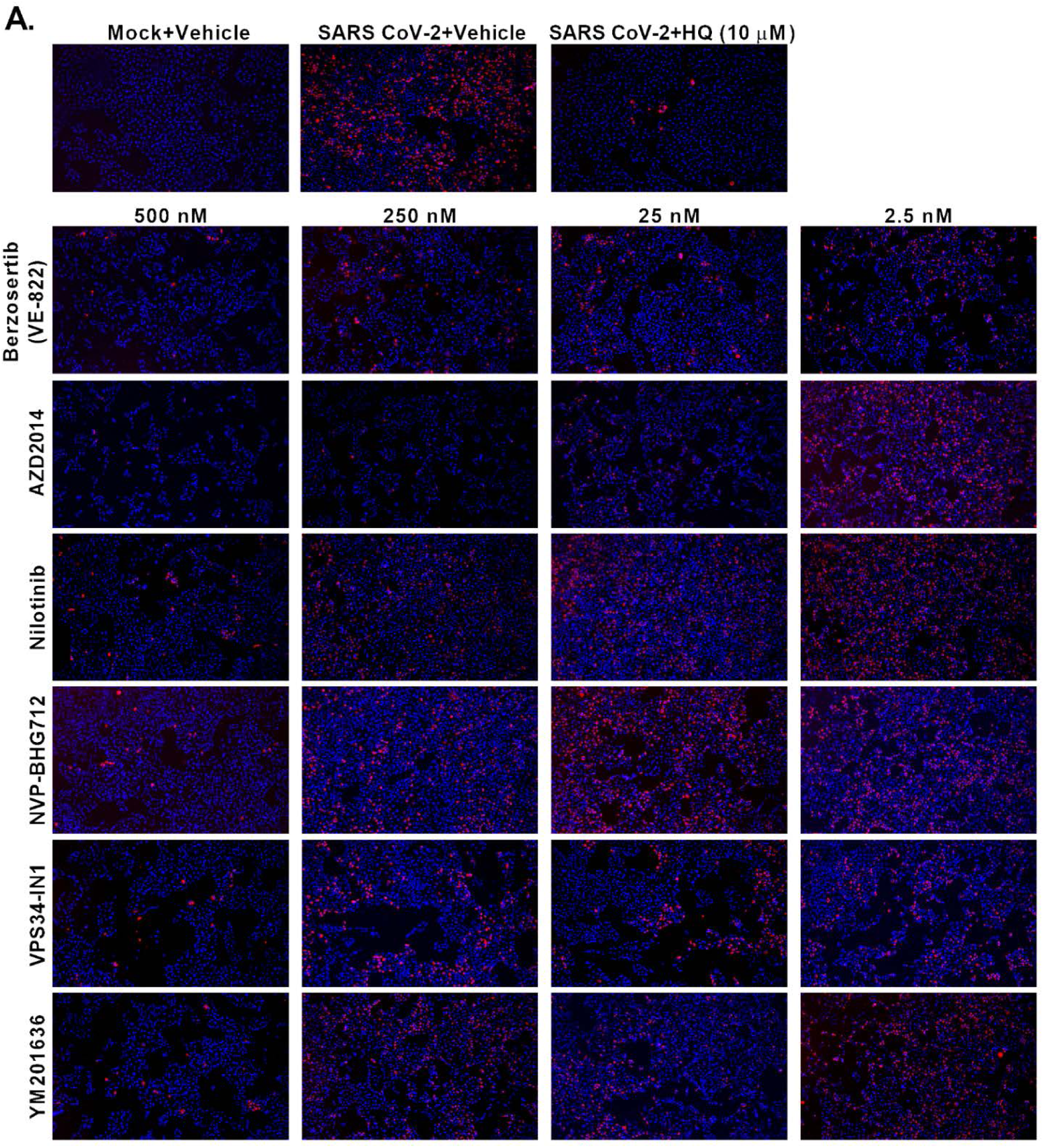
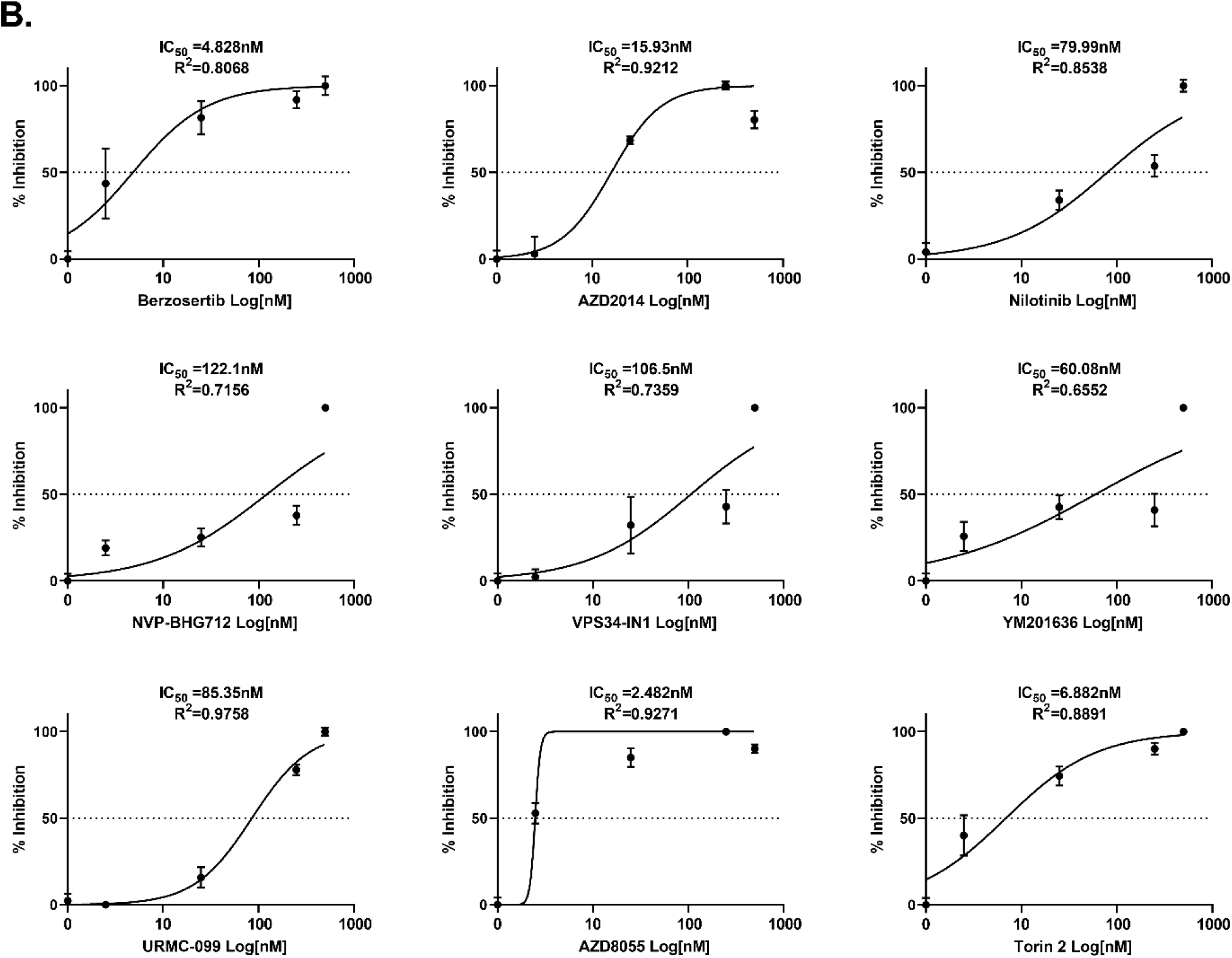
Secondary drug screen. (A) Immunofluorescent images of SARS-CoV-2 (red) infected cells treated with indicated drug compounds at various concentrations. (B) Graphs show percent inhibition of SARS-CoV-2 infectivity by indicated compounds. Note: IC50 of each compound is shown in the graph. Representative data from two independent experiments are presented.

### Secondary screen

From the primary screen, 34 compounds were selected for secondary screening with multiple drug doses (2.5, 25, 250 and 500 nM) in triplicates using 96-well plates. We used an immunofluorescent assay to quantify viral infected cells from each well. This comprehensive screen verified many hits of the primary screen (Figure 3A). Berzosertib (VE-822) and Vistusertib (AZD2014) demonstrated antiviral activities at IC50 of below 25 nM. Nilotinib, NVP-BHG712, VPS34-INI and YM201636 showed IC50 ranges between 50 nM – 125 nM (Figure 3B). These compounds act against SARS-CoV-2 by limiting viral infection through inhibiting critical cellular enzymes needed for viral replication. These kinase inhibitors are nucleoside analogues. Nucleoside analogues such as Remdesivir, EIDD-2801, and Ribavirin, have shown to inhibit viral RNA dependent RNA polymerase (RdRp) enzyme and inducing compromising errors during viral genome replication.

### Drug validation in human cells

The screens were conducted using Vero-E6, a monkey kidney epithelial cell line. Subsequently, we performed additional validation in human cells. We mainly focused on a new class of antiviral compound Berzosertib, which is already in Phase 2 clinical trial for cancer ailments. Initially, we have utilized an ACE2 entry receptor over expressing human embryonic kidney 293T cell line (293T-ACE2). The SARS-CoV-2 infected 293T-ACE2 cells were treated with Berzosertib (100 nM) and immunostaining analysis showed complete inhibition of virus replication at 48 hpi (Supplementary Figure 2). We also included hydroxychloroquine (10 µM) as a positive control. These observations provides additional confirmation of Berzosertib as a potential candidate for the treatment of SARS-CoV-2 infection.

COVID-19 patients have presented with cardiovascular complications. Recently, human induced pluripotent stem cell-derived cardiomyocytes (hiPSC-CM) have been shown to be great at recapitulating cardiovascular diseases at a cellular level^30,31^, and have demonstrated susceptibility to SARS-CoV-2 infection^32^. Thus this cardiomyocyte system would be useful for antiviral drug testing against SARS-CoV-2. To further evaluate the potency of Berzosertib, hiPSC-CMs were treated with this compound (250 nM) and the inhibition of virus production was evaluated by quantifying infectious virus in the supernatant at various timepoints (Figure 4A). We observed Berzosertib treatment had significantly reduced the virus production as well as apoptotic cell death associated with viral infection (Figure 4B-C). The infected cardiomyocytes were confirmed by specific staining of cardiac troponin T (cTnT) and viral Spike proteins (Figure 4D). Subsequently, untreated cells had infection-mediated cell injury with disrupted troponin T fibers. We specifically focused on toxicity associated with Berzosertib. We found treatment of cardiomyocytes infected SARS-CoV-2 with Berzosertib stabilized cardiomyocyte function with similar beats per minute to uninfected cardiomyocytes (Figure 4E, and Supplementary Videos). SARS-CoV-2 infected cells had significantly reduced cardiomyocyte beating with non-synchronous twitching of few cluster of cells (Refer to Supplementary Videos). Taken together, our results shows that Berzosertib is a potent, safe, and effective new class of antiviral agent against SARS-CoV-2.

**Figure 4.**
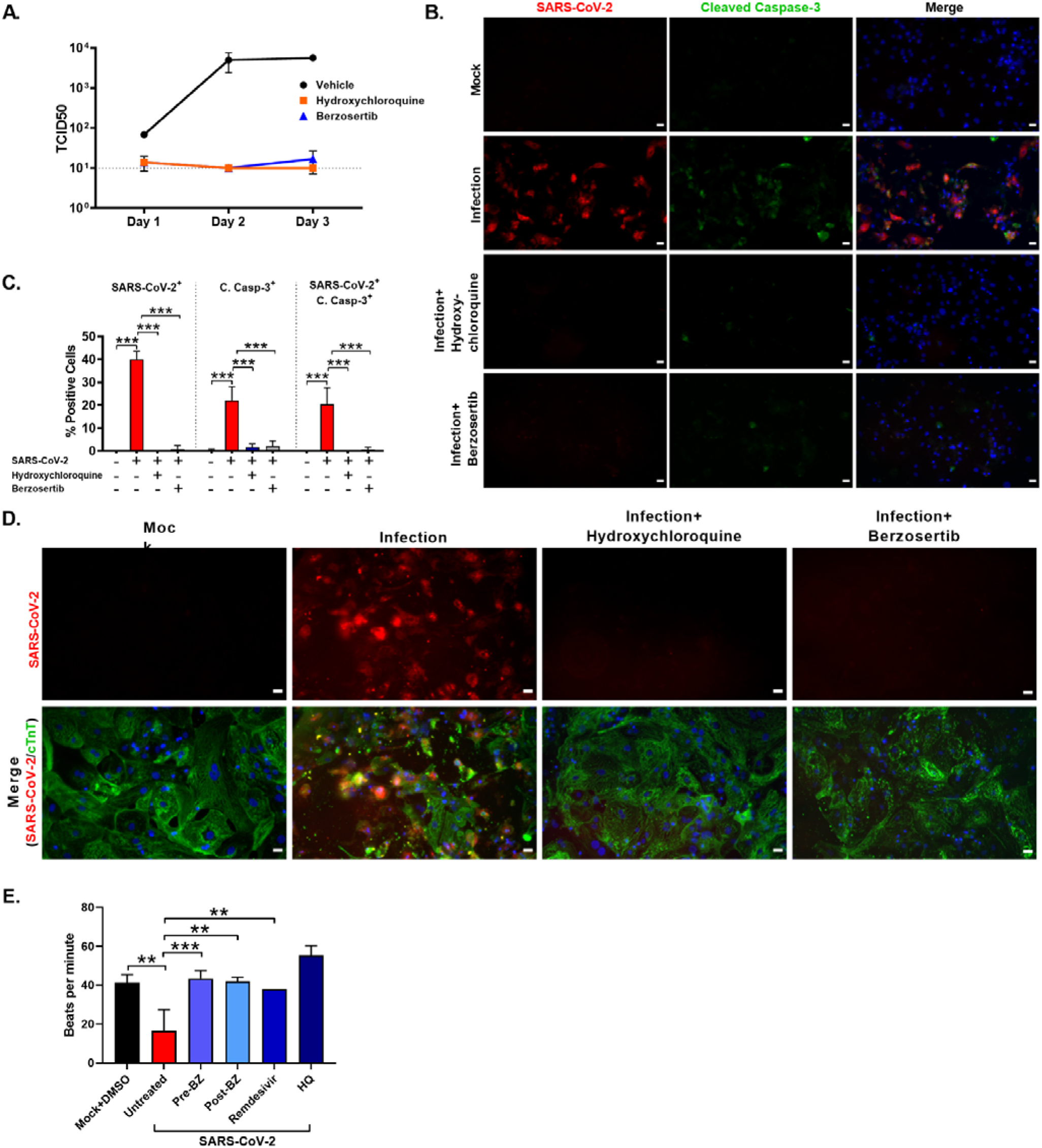
Berzosertib inhibits SARS-CoV-2 replication in human iPSC cardiomyocytes. (A) Graph shows viral titer (TCID50/25ul) of supernatant collected at indicated timepoints after SARS-CoV-2 infection of drug treated hiPSC-CMs. (B) IFA images of cells undergoing apoptosis assessed by cleaved caspase-3 staining on drug treated hiPSC-CMs at 72 hpi. Scale bar=25 µm. (C) Graph depicts quantification of SARS-CoV-2 and cleaved caspase-3 positive cells. (D) hiPSC-CMs stained for cardiac marker cardiac troponin T (cTnT) shows cells are protected from SARS-CoV-2 infection by drugs Berzosertib (250nM) and HQ (10µM). Scale bar=25 µm. (E). Graph shows beats per minute of SARS-CoV-2 infected hiPSC-CM cells treated with Berzosertib (250nM) (pre- and post -treatment), Remdesivir (10 µM), and HQ (10 µM). Statistical analysis of graphs (C, E): Multiple-comparison one-way analysis of variance (ANOVA) was conducted (***, P < 0.0001; **, P < 0.001). Representative data from two independent experiments are presented.

### Berzosertib’s Mode of Action

Berzosertib (Figure 5A) is a selective inhibitor of Serine/threonine-protein kinase ATR (ataxia telangiectasia and Rad3-related protein) which can inhibit the DNA Damage Response pathway. Previous studies have shown that Berzosertib blocks the phosphorylation of downstream signaling factor CHK1 (phospho-CHK1-S345) and potentiates phosphorylation of H2AX (phospho-H2AX-S139 or γH2AX) marker by DNA damaging agents^33,34^. It has been known that many viruses hijack this pathway for efficient replication^35,36^. One of the mode of actions could be inhibition of the aberrant activation of DNA repair signaling pathway by the virus, which can result in virus replication blockage. Indeed, we observed that SARS-CoV-2 infection has activated DDR pathway in Vero-E6 cells (Figure 5B). Currently, additional mechanistic and pre-clinical in vivo studies are being conducted for submission of an Investigational New Drug (IND) application towards clinical trials in COVID-19 patients.

**Figure 5.**
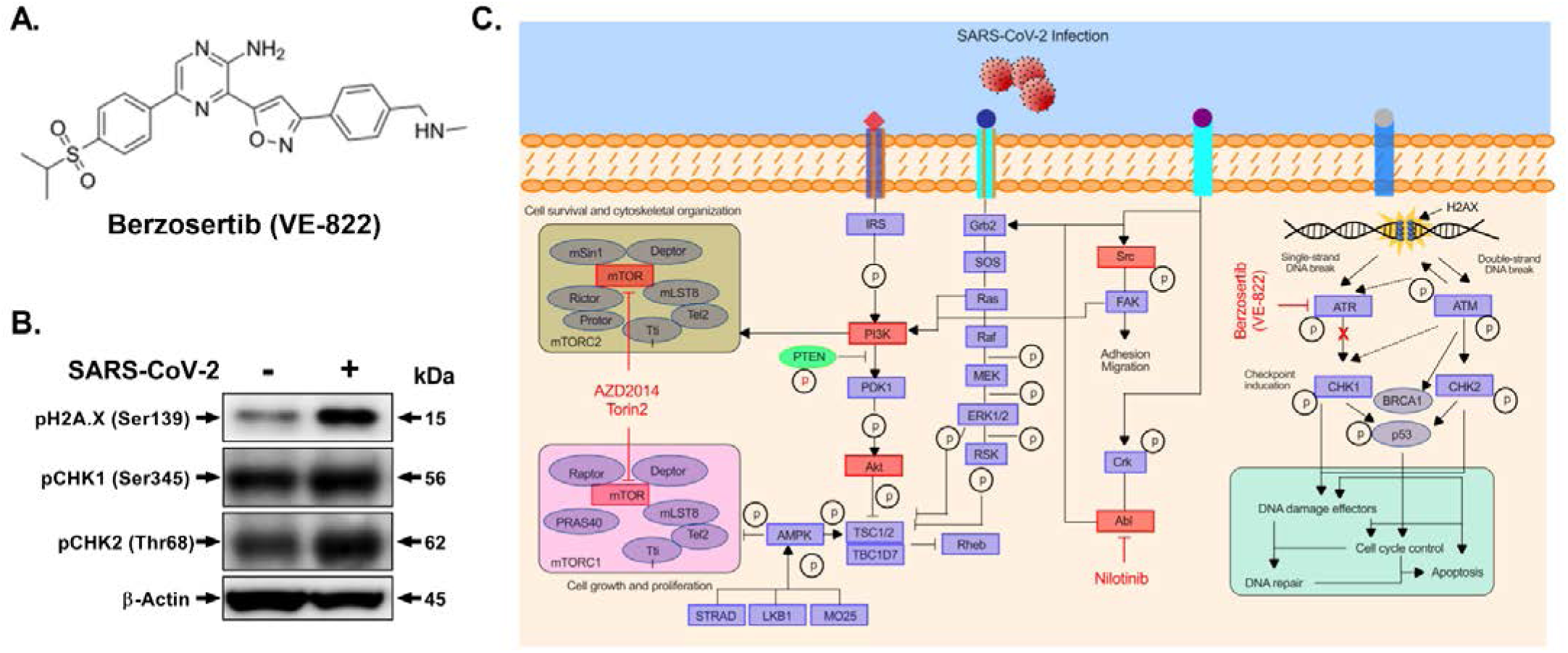
SARS-CoV-2 interaction with key cellular signaling pathways. (A) Chemical structure of compound Berzosertib is shown. (B) Western blot analysis shows phosphorylation of key DNA-Damage Response (DDR) proteins: CHK1, CHK2, and H2A.X. (C) Schematic illustration demonstrates the key pathways identified in the kinase inhibitor drug screen having critical role in SARS-CoV-2 infection. Drug compounds and their respective target kinase pathways (mTOR-PI3K-AKT, ABL-BCR/MAPK, and DNA-Damage Response) are described.

## DISCUSSION

We performed an antiviral drug screen using kinase inhibitors targeting SARS-CoV-2 infection. We have identified 34 drugs out of 430 compounds with anti-SARS-CoV-2 activity. These selected compounds mainly targeted mTOR-PI3K-AKT, ABL-BCR/MAPK, and DNA-Damage Response (DDR) Pathways (Figure 2B and 5C). Our secondary screen confirmed the antiviral activities of the following kinase inhibitors: AZD2014, Torin2, Nilotinib, NVP-BHG712, VPS34-INI, YM201636 and Berzosertib. Critical cellular pathways are subjected to activation, suppression, or some form of modulation by viral infection.

Interestingly, we have observed that several antiviral compounds targeted mTOR-PI3K-AKT pathway, including AZD2014 and Torin2 (Figure 2B). The mammalian target of rapamycin (mTOR) regulates cell growth, autophagy, and various metabolic processes^37,38^ and have been shown to be targeted by various viruses. Upon influenza virus infection, mTORC2 and PDPK1 (3-phosphoinositide-dependent protein kinase 1) differentially phosphorylate AKT. Viruses such as Kaposi’s Sarcoma Herpesvirus, Hepatitis C virus, Epstein-Barr virus, and Adenovirus are well known to activate PI3K ^37,39-42^. Torin compound has shown to inhibit virus replication by blocking mTOR kinase^43^. Our findings suggest that AZD2014 and Torin2 targeting PI3K/AKT1/MTOR pathway could be developed as potential therapeutics against COVID-19.

We identified a lead drug Berzosertib, which acts as an antiviral agent against SARS-CoV-2 by limiting viral genomic RNA replication. Berzosertib is a selective inhibitor of ATR kinase, which has been previously shown to block the DNA damage response pathway in cancer cells, with no discernable effect on normal cells ^44^. The DNA damage response pathway is responsible for maintaining cellular genome integrity which is also modulated by DNA tumor viruses^45^. An alphaherpesvirus, Marek’s disease virus (MDV), and BK polyomavirus (BKPyV) have been shown to induce the DDR during infection^46,47^. It is apparent that many RNA viruses can induce significant DNA damage, even in cases where viral replication takes place exclusively in the cytoplasm. DNA damage can contribute to the pathogenesis of RNA viruses through triggering of apoptosis, stimulation of inflammatory immune responses and the introduction of deleterious mutations that can increase the risk of tumorigenesis. In addition, activation of DDR pathways can contribute positively to replication of viral RNA genomes. Elucidation of the interactions between RNA viruses and the DNA damage response (DDR) would provide important insights into modulation of host cell functions by these pathogens.

For all of these reasons the DDR ATR signaling pathway appears to be a valid target for host-directed therapeutic development for COVID-19 and warrants further investigation. As of June 2020, there have been eight clinical studies (at Phase 1 and 2) based on Berzosertib and the side effect profile is mild and the drug is well tolerated. Further evaluation of the targets using CRISPR based KO studies is in progress. Thus, we consider this compound can be rapidly repurposed to treat COVID-19 patients. Overall, this present study illustrates key signaling proteins involved in SARS-CoV-2 replication and provides potential avenues for novel antiviral drug development.

## METHODS AND MATERIALS

### Ethics Statement

This study was performed in strict accordance with the recommendations of UCLA.

### Cell lines

Vero-E6 [VERO C1008 (ATCC® CRL-1586(tm))] cells were obtained from ATCC. Cells were cultured in EMEM growth media containing 10% fetal bovine serum (FBS) and penicillin (100 units/ml). ACE2 entry receptor overexpressing human embryonic kidney 293T cells (293T-ACE2) were established and cultured in the media described above with the presence of puromycin (1 µg/ml). Cells were incubated at 37°C with 5% CO_2_. The hiPSC-CMs were generated from hiPSCs by directed differentiation approach modulating Wnt signaling using a small-molecule^48^ and as previously described^48^, cardiomyocytes were metabolically selected by using glucose deprivation. After selection, hiPSC-CMs were replated for viral infection.

### Virus

SARS-Related Coronavirus 2 (SARS-CoV-2), Isolate USA-WA1/2020, was obtained from BEI Resources of National Institute of Allergy and Infectious Diseases (NIAID). All the studies involving live virus was conducted in UCLA BSL3 high-containment facility. SARS-CoV-2 was passaged once in Vero-E6 cells and viral stocks were aliquoted and stored at -80°C. Virus titer was measured in Vero-E6 cells by TCID_50_ assay.

### Biosafety and IRB Approval

Appropriate institutional review boards (IRB) approvals were obtained at UCLA and Cedars-Sinai Medical Center. All hiPSC lines used in this study have been approved by the UCLA and Cedars-Sinai Medical Center human pluripotent stem cell research oversight committees.

### SARS-CoV-2 Infection

Vero cells were seeded at 5 x 10^3^ cells per well in 0.2 ml volumes using a 96-well plate and hiPSC-CMs were replated at 1 x 10^5^ cells per well. The following day, viral inoculum (MOI of 0.01 and 0.1; 100 µl/well) was prepared using serum free media. The spent media from each well was removed and 100 µl of prepared inoculum was added onto Vero cells. For mock infection, serum free media (100 µl/well) alone was added. The inoculated plates were incubated for 1 hr at 37 °C with 5% CO_2_. The inoculum was spread by gently tilting the plate sideways at every 15 minutes. At the end of incubation, the inoculum was replaced with serum supplemented media (200 µl per well) and for hiPSC-CM, cell culture medium was replaced with RPMI 1640 + B27 supplement with insulin. At selected timepoints live cell images were obtained by bright field microscope. At 48 hours post infection (hpi), viral infection was examined by immunocytochemistry (ICC) analysis using SARS-CoV Spike (S) antibodies [BEI Resources: NR-10361 Polyclonal Anti-SARS Coronavirus (antiserum, Guinea Pig), and NR-616 Monoclonal Anti-SARS-CoV S Protein (Similar to 240C) SARS coronavirus].

### Antiviral Drug Study

Vero-E6 cells were seeded on 96-well plates and were pretreated with drugs for 1 hour, then SARS-CoV-2 inoculum (MOI 0.1) was added. DMSO vehicle treated cells, with or without viral infections, were included as controls. 48 hpi, the cells were fixed and immunostained with anti-dsRNA antibody (J2 clone; Absolute Antibody Inc, USA) to assess viral genome replication (Figure 1C).

### Immunohistochemistry

Cells were fixed with methanol (incubated in -20°C freezer until washed with PBS) or 4% Paraformaldehyde for 30-60 minutes. Cells were washed 3 times with 1x PBS and permeabilized using blocking buffer (0.3% Triton X-100, 2% BSA, 5% Goat Serum, 5% Donkey Serum in 1 X PBS) for 1 hour at room temperature. For immunostaining, cells were incubated overnight at 4°C with each primary antibody. The cells were then washed with 1X PBS three times and incubated with respective secondary antibody for 1 hour at room temperature. Nuclei were stained with DAPI (4’,6-Diamidino-2-Phenylindole, Dihydrochloride) (Life Technologies) at a dilution of 1:5000 in 1 X PBS. Image acquisition was done using Leica DM IRB fluorescent microscopes.

### Western Blot analysis

Cells were lysed in 50 mM Tris pH 7.4, 1% NP-40, 0.25% sodium deoxycholate, 1 mM EDTA, 150 mM NaCl, 1 mM Na3VO4, 20 Mm or NaF, 1mM PMSF, 2 mg ml^- 1^ aprotinin, 2 mg ml^-1^ leupeptin and 0.7 mg ml^-1^ pepstatin or Laemmli Sample Buffer (Bio Rad, Hercules, CA). Cell lysates were resolved by SDS-PAGE using 10% gradient gels and transferred to a 0.2 µm PVDF membrane. Subsequently, the membranes were blocked with 5% skim milk and 0.1% Tween-20 in 1x TBST (0.1% Tween-20) at room temperature for 1 hour. The membranes were then probed with respective monoclonal antibodies and detected by Thermo Scientific SuperSignal West Femto Maximum Sensitivity Substrate.

### Image Analysis/Quantification

Microscope images were obtained using the Leica DM IRB and Zeiss Software Program. Three to five images per well were quantified for each condition using Image J’s plugin Cell Counter feature was used to count the positively stained cells by a double blinded approach.

## Data analysis

All testing was done at the two-sided alpha level of 0.05. Data were analyzed for statistical significance using unpaired student’s *t*-test to compare two groups (uninfected vs. infected) with Graph Pad Prism software, version 8.1.2 (GraphPad Software, US).

## Supporting information

SARS-CoV-2 infected hiPSC-CM beats (3 dpi)

Berzosertib (250 nM) pre-treated and SARS-CoV-2 infected hiPSC-CM beats (3 dpi)

## Data availability

All relevant data regarding this manuscript is available from the above listed authors.

### Acknowledgments

This research was funded by UCLA DGSOM and Broad Stem Cell Research Center institutional award (OCRC #20-1) to V.A. and California Institute for Regenerative Medicine Discovery Award (DISC2COVID19-11764) to B.G and (TRAN1COVID19-11975) to V.A. The following reagents were obtained through BEI Resources, NIAID, NIH: Monoclonal Anti-SARS-CoV S Protein (Similar to 240C), NR-616; Polyclonal Anti-SARS Coronavirus (antiserum, Guinea Pig), NR-10361. The following reagent was deposited by the Centers for Disease Control and Prevention and obtained through BEI Resources, NIAID, NIH: SARS-Related Coronavirus 2, Isolate USA-WA1/2020, NR-52281. We thank David Austin for cell quantification at the Molecular Screening Shared Resource (MSSR) at UCLA. We thank Kouki Morizono for providing 293T-ACE2 cell line. We are grateful to Barbara Dillon, UCLA High Containment Program Director for BSL3 work.

## Author contributions

Garcia Jr. G: Conception and design, Collection and/or assembly of data. Data analysis and interpretation, and Manuscript writing. Sharma A., Sen C: Conducted experiments, Data analysis and interpretation. Ramaiah A., Kohn D., Gomperts B., Svendsen C.N., and Damoiseaux R.D.: Experimental design, Data analysis, interpretation and Manuscript writing. Arumugaswami V: Conception and design, Data analysis and interpretation, Manuscript writing and Final approval of manuscript.

## Competing interests

The authors declare no competing financial interests.

## Materials & Correspondence

Supplementary information is available for this paper. Correspondence and requests for materials should be addressed to Vaithilingaraja Arumugaswami.

## SUPPLEMENTARY MATERIALS

**Supplementary Figure 1.**
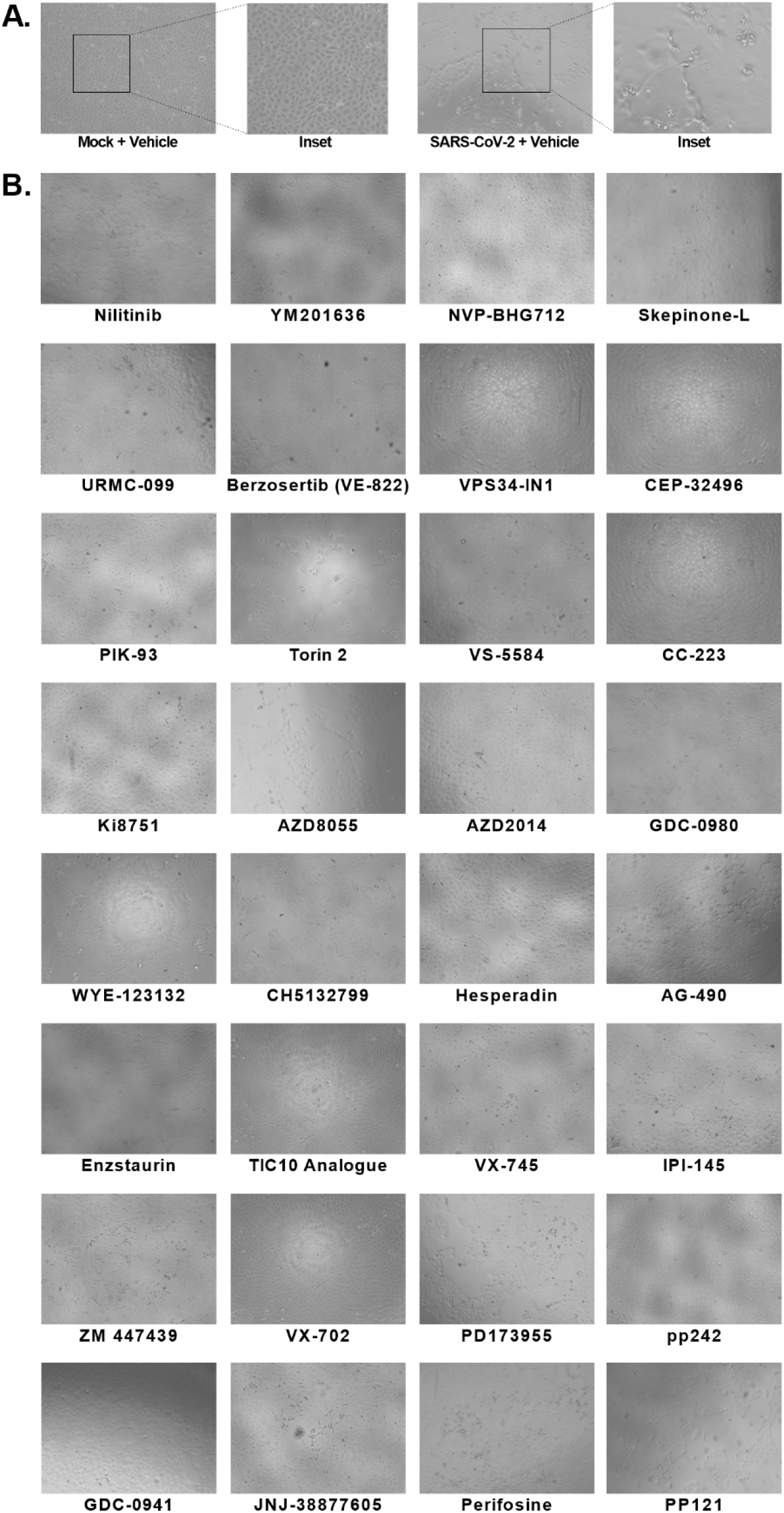
Primary screen of compounds inhibiting SARS-CoV-2 viral cytopathic effect. (A) DMSO Vehicle treated Vero-E6 cells had pronounced viral CPE at 48 hpi. Uninfected cells (Mock+vehicle) are included as negative control. (B) Bright field microscopic images (10X) of drug compounds treated SARS-CoV-2 infected cells showing no or reduced level of viral CPE.

**Supplementary Figure 2.**
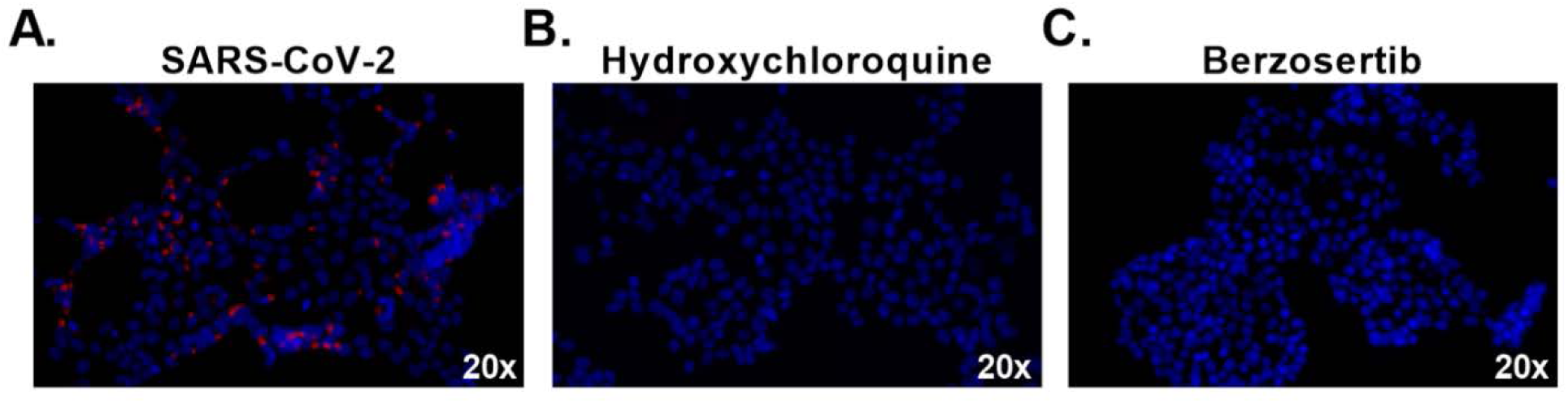
SARS-CoV-2 and Human 293T-ACE2 cell culture system for drug testing. IFA images show SARS-CoV-2 (Red) infection in untreated cells (A) and hydroxychloroquine (10 µM) treated cells (B). (C) Complete reduction of SARS-CoV-2 infection in Berzosertib (100nM) treated cells. 20x magnification.

**Supplementary Table 1**. List of 430 kinase inhibitors used in this study.

## Supplementary Videos

Supplementary Video 1. Mock hiPSC-CM beats (3 dpi).

Supplementary Video 2. SARS-CoV-2 infected hiPSC-CM beats (3 dpi).

Supplementary Video 3. Remdesivir (10µM) treated and SARS-CoV-2 infected hiPSC-CM beats (3 dpi).

Supplementary Video 4. Hydroxychloroquine (10µM) treated and SARS-CoV-2 infected hiPSC-CM beats (3 dpi).

Supplementary Video 5. Berzosertib (250 nM) pre-treated and SARS-CoV-2 infected hiPSC-CM beats (3 dpi).

Supplementary Video 6. Berzosertib (250 nM) post-treated and SARS-CoV-2 infected hiPSC-CM beats (3 dpi).

## Notes

### Competing Interest Statement

The authors have declared no competing interest.

## REFERENCES

1. Dong E, Du H, Gardner L. An interactive web-based dashboard to track COVID-19 in real time. The Lancet Infectious Diseases. 2020;20(5):533–534.

2. Worldometers.info. 2020; https://www.worldometers.info/faq/. Accessed 23 June, 2020.

3. Ramaiah A, Arumugaswami V. Insights into Cross-species Evolution of Novel Human Coronavirus 2019-nCoV and Defining Immune Determinants for Vaccine Development. bioRxiv. 2020:2020.2001.2029.925867.

4. Kai H, Kai M. Interactions of coronaviruses with ACE2, angiotensin II, and RAS inhibitors-lessons from available evidence and insights into COVID-19. Hypertension research : official journal of the Japanese Society of Hypertension. 2020;43(7):648-654. PMC7184165

5. Hou YJ, Okuda K, Edwards CE, et al. SARS-CoV-2 Reverse Genetics Reveals a Variable Infection Gradient in the Respiratory Tract. Cell. 2020.

6. Xu Z, Shi L, Wang Y, et al. Pathological findings of COVID-19 associated with acute respiratory distress syndrome. The Lancet Respiratory Medicine. 2020;8(4):420–422.

7. Pacciarini F, Ghezzi S, Canducci F, et al. Persistent Replication of Severe Acute Respiratory Syndrome Coronavirus in Human Tubular Kidney Cells Selects for Adaptive Mutations in the Membrane Protein. Journal of virology. 2008;82(11):5137–5144.

8. Fanelli V, Fiorentino M, Cantaluppi V, et al. Acute kidney injury in SARS-CoV-2 infected patients. Critical Care. 2020;24(1):155.

9. Puelles VG, Lütgehetmann M, Lindenmeyer MT, et al. Multiorgan and Renal Tropism of SARS-CoV-2. New England Journal of Medicine. 2020.

10. Varga Z, Flammer AJ, Steiger P, et al. Endothelial cell infection and endotheliitis in COVID-19. Lancet (London, England). 2020;395(10234):1417-1418. PMC7172722

11. Shi S, Qin M, Shen B, et al. Association of Cardiac Injury With Mortality in Hospitalized Patients With COVID-19 in Wuhan, China. JAMA Cardiology. 2020.

12. Fried JA, Ramasubbu K, Bhatt R, et al. The Variety of Cardiovascular Presentations of COVID-19. Circulation. 2020;141(23):1930–1936.

13. Dobrovolny HM, Beauchemin CAA. Modelling the emergence of influenza drug resistance: The roles of surface proteins, the immune response and antiviral mechanisms. PLoS One. 2017;12(7):e0180582. PMC5503263

14. Bright RA, Shay DK, Shu B, Cox NJ, Klimov AI. Adamantane resistance among influenza A viruses isolated early during the 2005-2006 influenza season in the United States. Jama. 2006;295(8):891–894.

15. Supekova L, Supek F, Lee J, et al. Identification of human kinases involved in hepatitis C virus replication by small interference RNA library screening. The Journal of biological chemistry. 2008;283(1):29–36.

16. Li Q, Brass AL, Ng A, et al. A genome-wide genetic screen for host factors required for hepatitis C virus propagation. Proceedings of the National Academy of Sciences of the United States of America. 2009;106(38):16410-16415. PMC2752535

17. Keating JA, Striker R. Phosphorylation events during viral infections provide potential therapeutic targets. Reviews in medical virology. 2012;22(3):166-181. PMC3334462

18. Jiang WM, Zhang XY, Zhang YZ, Liu L, Lu HZ. A high throughput RNAi screen reveals determinants of HIV-1 activity in host kinases. International journal of clinical and experimental pathology. 2014;7(5):2229-2237. PMC4069921

19. Gross S, Rahal R, Stransky N, Lengauer C, Hoeflich KP. Targeting cancer with kinase inhibitors. The Journal of clinical investigation. 2015;125(5):1780-1789. PMC4463189

20. Ott PA, Adams S. Small-molecule protein kinase inhibitors and their effects on the immune system: implications for cancer treatment. Immunotherapy. 2011;3(2):213-227. PMC4009988

21. Development TCftSoD. Cost to develop and win 279 marketing approval for a new drug is $2.6 billion. 2014; http://csdd.tufts.edu/news/complete_story/pr_tufts_csdd_2014_cost_study.

22. Schor S, Einav S. Repurposing of Kinase Inhibitors as Broad-Spectrum Antiviral Drugs. DNA Cell Biol. 2018;37(2):63–69.

23. García M, Cooper A, Shi W, et al. Productive replication of Ebola virus is regulated by the c-Abl1 tyrosine kinase. Sci Transl Med. 2012;4(123):123ra124–123ra124.

24. Johansen LM, Brannan JM, Delos SE, et al. FDA-approved selective estrogen receptor modulators inhibit Ebola virus infection. Sci Transl Med. 2013;5(190):190ra179–190ra179.

25. Madrid PB, Chopra S, Manger ID, et al. A systematic screen of FDA-approved drugs for inhibitors of biological threat agents. PLoS One. 2013;8(4):e60579–e60579.

26. Riva L, Yuan S, Yin X, et al. A Large-scale Drug Repositioning Survey for SARS-CoV-2 Antivirals. bioRxiv. 2020:2020.2004.2016.044016.

27. de Wilde AH, Jochmans D, Posthuma CC, et al. Screening of an FDA-approved compound library identifies four small-molecule inhibitors of Middle East respiratory syndrome coronavirus replication in cell culture. Antimicrob Agents Chemother. 2014;58(8):4875–4884.

28. Liu J, Cao R, Xu M, et al. Hydroxychloroquine, a less toxic derivative of chloroquine, is effective in inhibiting SARS-CoV-2 infection in vitro. Cell discovery. 2020;6:16. PMC7078228

29. Kuhn M, von Mering C, Campillos M, Jensen LJ, Bork P. STITCH: interaction networks of chemicals and proteins. Nucleic acids research. 2008;36(Database issue):D684-688. PMC2238848

30. Lan F, Lee AS, Liang P, et al. Abnormal calcium handling properties underlie familial hypertrophic cardiomyopathy pathology in patient-specific induced pluripotent stem cells. Cell stem cell. 2013;12(1):101-113. PMC3638033

31. Sun N, Yazawa M, Liu J, et al. Patient-specific induced pluripotent stem cells as a model for familial dilated cardiomyopathy. Sci Transl Med. 2012;4(130):130ra147–130ra147.

32. Sharma A, Garcia G, Arumugaswami V, Svendsen CN. Human iPSC-Derived Cardiomyocytes are Susceptible to SARS-CoV-2 Infection. Cell Reports Medicine. 2020.

33. Hall AB, Newsome D, Wang Y, et al. Potentiation of tumor responses to DNA damaging therapy by the selective ATR inhibitor VX-970. Oncotarget. 2014;5(14):5674-5685. PMC4170644

34. Sanjiv K, Hagenkort A, Calderon-Montano JM, et al. Cancer-Specific Synthetic Lethality between ATR and CHK1 Kinase Activities. Cell reports. 2016;14(2):298-309. PMC4713868

35. Boichuk S, Hu L, Hein J, Gjoerup OV. Multiple DNA damage signaling and repair pathways deregulated by simian virus 40 large T antigen. Journal of virology. 2010;84(16):8007-8020. PMC2916509

36. Lilley CE, Schwartz RA, Weitzman MD. Using or abusing: viruses and the cellular DNA damage response. Trends in microbiology. 2007;15(3):119–126.

37. Le Sage V, Cinti A, Amorim R, Mouland AJ. Adapting the Stress Response: Viral Subversion of the mTOR Signaling Pathway. Viruses. 2016;8(6):152.

38. Weichhart T. Mammalian target of rapamycin: a signaling kinase for every aspect of cellular life. Methods in molecular biology (Clifton, NJ). 2012;821:1–14.

39. Sodhi A, Chaisuparat R, Hu J, et al. The TSC2/mTOR pathway drives endothelial cell transformation induced by the Kaposi’s sarcoma-associated herpesvirus G protein-coupled receptor. Cancer cell. 2006;10(2):133–143.

40. Bose SK, Shrivastava S, Meyer K, Ray RB, Ray R. Hepatitis C virus activates the mTOR/S6K1 signaling pathway in inhibiting IRS-1 function for insulin resistance. Journal of virology. 2012;86(11):6315-6322. PMC3372214

41. Kong K, Kumar M, Taruishi M, Javier RT. The human adenovirus E4-ORF1 protein subverts discs large 1 to mediate membrane recruitment and dysregulation of phosphatidylinositol 3-kinase. PLoS pathogens. 2014;10(5):e1004102. PMC4006922

42. Moody CA, Scott RS, Amirghahari N, et al. Modulation of the cell growth regulator mTOR by Epstein-Barr virus-encoded LMP2A. Journal of virology. 2005;79(9):5499-5506. PMC1082717

43. Kuss-Duerkop SK, Wang J, Mena I, et al. Influenza virus differentially activates mTORC1 and mTORC2 signaling to maximize late stage replication. PLoS pathogens. 2017;13(9):e1006635. PMC5617226

44. Mei L, Zhang J, He K, Zhang J. Ataxia telangiectasia and Rad3-related inhibitors and cancer therapy: where we stand. Journal of hematology & oncology. 2019;12(1):43. PMC6482552

45. Pancholi NJ, Price AM, Weitzman MD. Take your PIKK: tumour viruses and DNA damage response pathways. Philosophical transactions of the Royal Society of London Series B, Biological sciences. 2017;372(1732). PMC5597736

46. Lian X, Bao C, Li X, et al. Marek’s Disease Virus Disables the ATR-Chk1 Pathway by Activating STAT3. Journal of virology. 2019;93(9). PMC6475774

47. Verhalen B, Justice JL, Imperiale MJ, Jiang M. Viral DNA replication-dependent DNA damage response activation during BK polyomavirus infection. Journal of virology. 2015;89(9):5032-5039. PMC4403456

48. Sharma A, Li G, Rajarajan K, Hamaguchi R, Burridge PW, Wu SM. Derivation of highly purified cardiomyocytes from human induced pluripotent stem cells using small molecule-modulated differentiation and subsequent glucose starvation. Journal of visualized experiments : JoVE. 2015(97). 4401368

